# Transposable Elements Profiling Reveals DUXA-associated MLT1D Endogenous Retroviral Elements Activation During Bovine Maternal to Zygotic Transition

**DOI:** 10.64898/2026.09.23.753920

**Authors:** Guangsheng Li, Cédric Feschotte, Jingyue (Ellie) Duan

## Abstract

Transposable elements (TEs) are a major source of genomic diversity in mammals, yet their regulatory roles in the bovine genome remain poorly understood. Through characterizing bovine TE landscape, despite the substantial proportion (25.6%) of ruminant-specific TEs, we observe age- and class-dependent genomic distribution patterns similar to those observed in other mammals. Next, we profile TE and gene expression dynamics in pre-implantation embryos generated *in vivo* (IVV), by *in vitro* fertilization (IVT), and through somatic cell nuclear transfer (SCNT). The zygotic genome activation (ZGA) is shifted from the 4-cell stage to the 8-cell stage in IVT and SCNT embryos compared to IVV embryos. SCNT embryos exhibit impaired initiation of early transcription programs at the 4-cell stage and disrupted developmental trajectories, including abnormal activation of pluripotency-associated genes. A subset of retroviral LTR elements are strongly activated at ZGA in IVV and IVT embryos, whereas their activation is markedly muted in SCNT embryos, suggesting that impaired gene and TE reprogramming may contribute to the developmental defects commonly observed in SCNT embryos. By epigenomic profiling, the MLT1D elements from the ERVL-MaLR LTR family lose repressive marks and gain H3K27ac at ZGA, together with DUXA*-*binding motif enrichment. Knockdown of DUXA in bovine embryos significantly reduced MLT1D expression and ZGA marker genes. We propose that a subset of DUXA*-*enriched MLT1D functions as enhancers that promote ZGA. Overall, our study provides new insights into the regulatory roles of TEs during bovine embryogenesis and establishes a framework for comparative studies of TE-mediated gene regulation in early mammalian development.

## INTRODUCTION

Transposable elements (TEs) are repetitive DNA sequences capable of moving within the genome and comprising a substantial proportion of mammalian genomes (Osmanski et al., 2023; Wicker et al., 2007). TEs are broadly classified into DNA transposons, which follow the “cut and paste” mechanism mediated by transposase, and retrotransposons, which exhibit the “copy and paste” strategy involving reverse transcription (Lanciano and Cristofari, 2020). In humans, retrotransposons take up more than 40% of the genome, with long interspersed elements (LINEs) being the most prevalent TE class (Hoyt et al., 2022), whereas DNA transposons are generally less than 5% of the mammalian genomes (Osmanski et al., 2023). Among livestock species genomes, including chicken, pig, and cattle, TE composition also differs significantly. Approximately 10% of the chicken genome is composed of TEs, whereas pig and cattle genomes each harbor more than 40%, with LINEs being the most abundant retrotransposon class in all three species (Wang et al., 2025).

Because TEs have been continuously active throughout mammalian evolution, each species has a unique TE assortment with a mixture of ancestral elements (shared with distant mammals) and lineage-specific or even species-specific elements (Osmanski 2023). In the bovine genome, more than half of TEs are ruminant-specific, meaning they have inserted in the past 50 million years (Chen et al., 2019). Notably, multiple waves of LINEs, short interspersed elements (SINEs) and endogenous retroviruses (ERVs) expansion have occurred in the ruminant lineage (Wang et al., 2025; Zhao et al., 2026). Diverse TEs appeared to have retained transposition activity in the modern bovine genome, including two LINE superfamilies, RTE-BovB and L1, the Core-RTE SINEs (mobilized by BovB LINEs), and two ERV superfamilies, ERVK and ERV1 (Kelly et al., 2022; Tang et al., 2024; Wang et al., 2025; Zhao et al., 2026). By contrast, in humans only L1 LINEs and SINEs mobilized by L1 (e.g. Alu and SVA) are currently active (Huang et al., 2012). In particular, ERVK superfamilies have contributed substantially to TE expansion within the bovidae lineage, including the activity of eight bovine-specific ERVKs (Zhao et al., 2026). Together, these findings indicate that the bovine genome harbors a distinct TE landscape compared to more extensively studied mammals such as human and mouse.

Although TE mobility is often considered disruptive to genome integrity, the accumulation of TEs in mammalian genomes has provided abundant material fueling evolutionary novelties shaping genomic evolution, chromatin organization, and transcriptional regulation (DiRusso and Clark, 2023; Fueyo et al., 2022; Lawson et al., 2023). Notably, during preimplantation embryonic development, ERVs and other retroelements undergo widespread transcriptional activation in species-specific manner while serving potentially conserved functions in zygotic genome activation (ZGA) and cell fate regulation (Fu et al., 2019; Grow et al., 2015; Guo et al., 2024; Li et al., 2026; Solberg et al., 2024). In mice, MERVL elements are robustly activated at the 2-cell stage and promote totipotency- and ZGA-associated transcription activation (Peaston et al., 2004). MERVL elements also generate chimeric transcripts with more than 300 ZGA genes (Macfarlan et al., 2012), and further studies reveal that abnormal expression of MERVL elements potentially leads to developmental arrest at the 2-cell stage and embryonic lethality (Huang et al., 2017; Sakashita et al., 2023). In humans, a subset of the ERV elements, HERVL, is transcriptionally activated during ZGA and can be regulated by the DUX4 transcription factor, which is belong to the DUX family that exhibits conserved roles in activating ERVs upon mouse ZGA (De Iaco et al., 2017; Göke et al., 2015; Hendrickson et al., 2017; Pontis et al., 2019; Whiddon et al., 2017). HERVK and HERVH can function as long-range enhancers regulating transcription of coding and non-coding genes (Gerdes et al., 2016; Grow et al., 2015; Pontis et al., 2019), while human-specific HERVK LTR5Hs act as epiblast-specific enhancers required for blastocyst development (Fueyo et al., 2025). Moreover, HERVL-associated MLT2A1 LTRs were shown to promote human ZGA by forming chimeric zygotic transcripts and recruiting HNRNPU and RNA Pol II to regulate transcription in trans (Xiang et al., 2026). Together, these studies suggest a striking convergence in mouse and human in which young, lineage-specific ERVs have been repeatedly co-opted as regulatory elements during mammalian pre-implantation development.

Dairy cattle are economically important livestock, and assisted reproductive technologies such as in vitro fertilization (IVF) and embryo transfer are widely used to generate elite offspring (Nowicki, 2021; Peterson and Mitloehner, 2021). However, IVF embryos exhibit reduced developmental competence compared with in vivo-derived embryos, resulting in lower pregnancy success and high rates of early embryonic loss (Ealy et al., 2019). Although gene expression and epigenome dynamics during bovine ZGA have been characterized (Graf et al., 2014; Hu, B. et al., 2024; Jiang et al., 2014), only a few studies focus on TE expression during bovine embryogenesis. Particularly, several bovine ERV families, including ERV1-1_BT, BERV-K1, and BERV-K2 elements, are known to be activated during ZGA and placenta development, and the ERV expression levels are associated with cellular pluripotency (Baba et al., 2011; Bui et al., 2009; Khazaee et al., 2018). Moreover, recent cross-species analyses of mammalian embryos also revealed widespread TE-driven transcripts and bovine-specific TE activation during ZGA (Oomen et al., 2025). However, despite these advances, the regulatory role of TEs in bovine ZGA remains poorly characterized. To address this gap, we performed a comprehensive genomic and transcriptomic analysis of bovine TEs and profiled their expression dynamics during preimplantation development across embryos generated under different conditions. This study provides a foundation for understanding TE-mediated transcriptional regulation during bovine embryogenesis.

## RESULTS

### Ruminant-specific TE shapes the bovine genome

According to Repeatmasker annotation of the latest bovine genome assembly (ARS-UCD 2.0, 2023) TEs account for 46.29% of the bovine genome, including 2.15% of DNA transposons and 44.1% retrotransposons (**Supplemental Fig. S1A**). As in other mammals, LINEs are the most abundant retrotransposons, occupying around about 30% of the genome, whereas LTRs/ERVs account for less than 5% and are primarily composed of elements from the ERV1 and ERVL-MaLR superfamilies (**Supplemental Fig. S1A**). We further classified bovine TEs into two main categories based on the TE phylogeny of Dfam database, where TEs present in the ancestors of ruminant lineage are defined as ancestral TEs and the other TEs are treated as ruminant TEs (**Fig. 1A-1B and Supplemental Table S1,** see Methods). Ruminant TEs accounted for nearly two-thirds of the total TE genomic coverage (or ∼25% of the bovine genome), and as expected, exhibited significantly lower sequence divergence to their consensus sequences, consistent with more recent amplification (**Fig. 1A-1B**). We note that ruminant-specific LINEs contribute considerably greater genomic coverage than ancestral LINEs (17.24% vs 9.75%), likely due to their longer insertion length (**Supplemental Fig. S1B-1C**). At superfamily level, ruminant-specific TEs were dominated by the RTE-BovB LINE clade and tRNA-Core-RTE SINE clade (**Fig. 1C**), consistent with their extensive propagation within ruminant genomes (Ivancevic et al., 2018; Kordiš and Gubenšek, 1999). Importantly, both groups of elements are completely absent from the human and mouse genome because RTE-Bov-B LINEs were introduced by horizontal transfer from snakes into a ruminant ancestor (Ivancevic et al., 2018; Kordis and Gubensek, 1998, 1999) and Core-RTE SINEs were subsequently derived from them (Kojima, 2018).

**Fig. 1.**
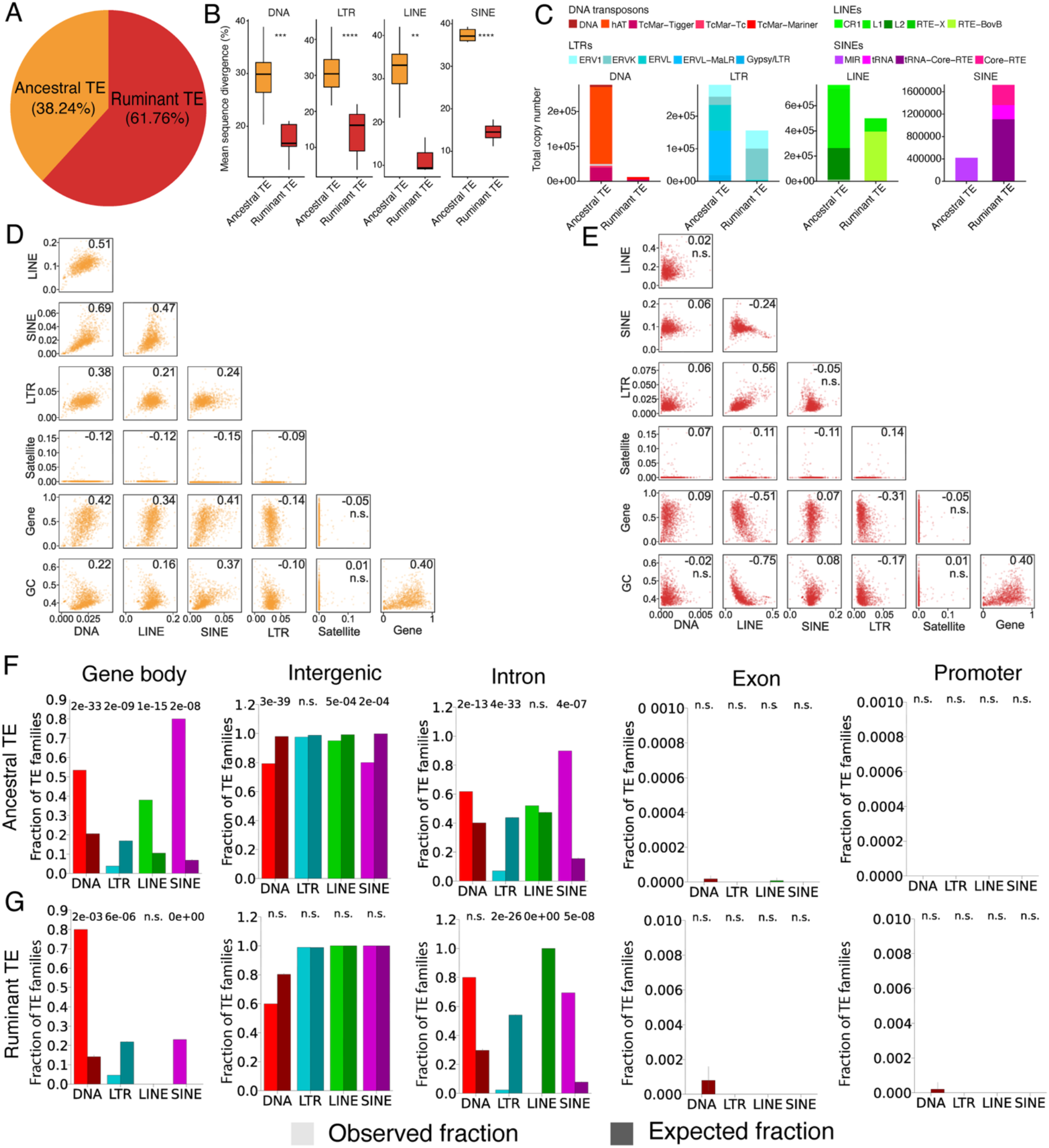
Genomic features of bovine TE. A: The relative proportions of ancestral and ruminant TE in bovine genome. B: Kimura sequence divergence comparison between ancestral and ruminant TE. C: Total copy number of ancestral and ruminant TE across different classes. The proportions of the main TE superfamilies are highlighted in the bar plot. D: Scatter plots show the Spearman rank correlation of genomic distribution between ancestral TE classes, satellite marks, GC content and genes. The correlation coefficients are shown on the top of each plot, and n.s. indicates no significance. E: Scatter plots show the Spearman rank correlation of genomic distribution between ruminant TE classes, satellite marks, GC content and genes. The correlation coefficients are shown on the top of each plot, and n.s. indicates no significance. F: Genomic distribution of ancestral TE in different functional regions. Within each TE class, the left bar indicates the observed fractions of TE family overlapped with the corresponding regions and the right bar indicated the expected fractions of TE family overlapped with the corresponding regions. The error bar of expected fraction indicates the 95% confidence interval of the mean value. The p-value is computed with binomial tests. G: Genomic distribution of ruminant TE in different functional regions. Within each TE class, the left bar indicates the observed fractions of TE family overlapped with the corresponding regions and the right bar indicated the expected fractions of TE family overlapped with the corresponding regions. The error bar of expected fraction indicates the 95% confidence interval of the mean value. The p-value is computed with binomial tests.

### Recent expansion of ruminant-specific retrotransposons

To more finely investigate the evolutionary history of bovine TEs, we estimated their age by phylogenetic analysis of 677 TE families with at least 10 copies and defragmented insertion length >= 100bp. For each family, median terminal branch length (substitutions per site) was calculated, where longer branches indicate a greater sequence divergence and older age, whereas shorter branches reflect more recent evolutionary activity, thus younger (**Supplemental Fig. S1D**). This method minimizes potential biases from family substructure (Chang et al., 2022; Stitzer et al., 2021). The resulting age estimates were consistent (Spearman’s ρ =0.55, *P* = 8.34E-55) with those based on mean divergence from family consensus sequences (**Supplemental Fig. S1E**).

DNA transposons were generally older (median branch length >0.55), including hAT-5_Mam, Charlie1a/4a, MER46C/103C, Arthur2, Zaphod4a, and Tigger13a, consistent with their amplification before placental mammal divergence (Chinwalla et al., 2002; Pace and Feschotte, 2007). Younger TE families (median branch length <0.23) were characterized by bovine-specific LINE and SINE families L1_BT and L1-3_BT, SINE2-1_BT and SINE2-3_BT, but the youngest families (median branch length <0.05, **Supplemental Table S2**) belong to the ERVK superfamily, including BTLTR1B/1C, BTLTR1, ERV2-1, and ERV2-3, suggesting very recent and possibly ongoing ERV transpositional activity in the bovine genome. Across all TE classes, family copy number showed a moderate positive correlation with evolutionary age (Spearman’s ρ = 0.23, P = 1.60E-09), a trend primarily driven by DNA transposons and LTRs (**Supplemental Fig. S1D**). However, only a few evolutionarily young (median branch length <0.13) TE families display extremely high copy number (>200,000 copies), including the SINE families BOV-A2, Bov-tA1/2/3, and BovB from LINEs (**Supplemental Table S3**). Collectively, these results reveal distinct evolutionary histories among bovine TE classes, characterized by ancient DNA transposons, lineage-specific expansion of LINEs and SINEs, and recent diversification of ERVs.

### Ruminant-specific TEs exhibit distinct genomic distribution patterns

Previous studies in human, mouse, and rat genomes have shown that TE distribution is not uniform along chromosomes and is strongly influenced by TE class and evolutionary age, GC content, and gene density (Chinwalla et al., 2002; Gibbs et al., 2004; Kvikstad and Makova, 2010; Lander et al., 2001; Medstrand et al., 2002; Soriano et al., 1983). In the bovine genome, we found that LINEs and LTRs exhibited colocalized enrichment in gene-poor, GC-low, and putative pericentromeric regions with satellite DNA markers, whereas SINEs and DNA transposons were enriched in gene-rich and GC-rich regions (**Supplemental Fig. S2A-B**). Furthermore, LINEs and LTRs were positively correlated (Spearman’s ρ = 0.44) with each other but negatively correlated (Spearman’s ρ ranging from -0.11 to -0.46) with SINEs and DNA transposons. Those TE class-specific patterns are consistent with those previously reported in other mammalian genomes (Correa et al., 2021; Kosovsky et al., 2025; Wang et al., 2025), suggesting broadly conserved mammalian TE partitioning patterns.

However, when separating ancestral and ruminant-specific TEs, we observe distinct distribution patterns unique to the bovine genome (**Fig. 1D-1E**). Interestingly, all ancestral TEs were positively colocalized (Spearman’s ρ = 0.21 - 0.69), with ancestral DNA transposons, LINEs, and SINEs showing positive correlations with gene dense and GC-rich regions (**Fig. 1D**). In contrast, unlike ancestral TEs, ruminant-specific LINEs and LTRs exhibited strengthened colocalization (Spearman’s ρ = 0.56), while ruminant LINEs were negatively correlated (Spearman’s ρ = -0.24) with ruminant-specific SINEs (**Fig. 1E**). Notably, ruminant LINEs, LTRs, and DNA transposons shifted toward gene-poor and low GC regions, while ruminant SINEs displayed a more uniform genomic distribution (**Fig. 1E**). Collectively, these findings indicate that the overall TE genomic distribution patterns are broadly conserved across mammals, while ruminant-specific TEs exhibit distinct genomic organization within the bovine genome.

### Distinct intergenic enrichment of ruminant-specific TEs

The genomic distributions of TEs are shaped by both insertion site preference and evolutionary selection against deleterious effects (Blass et al., 2012; Boissinot et al., 2006; Petrov et al., 2003). To examine TE localization in functional genomic regions, we analyzed TE enrichment across gene bodies, intergenic regions, introns, exons, and promoters (**Supplemental Fig. S2C,** see methods). Consistent with previous studies in mammalian genomes (Medstrand et al., 2002), all TE classes were predominantly enriched in intergenic regions instead of gene body regions. Within gene bodies, DNA transposons and SINEs were significantly enriched in introns compared to the expectation. In contrast, LTRs were significantly underrepresented in introns. Particularly, there were no TE families showing significantly more enrichments in exons or promoters than the expectation (**Supplemental Fig. S2C**). These patterns are similar to those observed in other organisms (Chang et al., 2022), indicating that TE distributions are shaped by both structural constraints, such as TE insertion length, and selective pressures imposed by gene-rich regions.

When separating ancestral and ruminant-specific TEs, distinct bovine-specific patterns emerged (**Fig. 1F-1G**). Notably, ∼20% of ruminant SINE families showed preferential location in gene body regions, whereas near 80% of ancestral SINE families were classified as preferentially intragenic, suggesting SINEs shift toward gene-rich regions as they get older. Comparing to ancestral DNA transposons, ruminant DNA transposons exhibited slightly increase in the fraction of preferentially intragenic and intronic families. For LTRs, both ruminant and ancestral elements maintained similar distributions across different genomic regions. Most strikingly, the three ruminant LINE families were all determined as preferentially intergenic distributions, while ∼ 35% of ancestral LINE families were shown as preferentially intragenic localizations. To investigate the selective forces underlying the preferential intergenic enrichment of ruminant LINEs, we analyzed the three ruminant LINE families and found that BovB accounted for about 75% of total ruminant LINE coverage, followed by L1_BT and L1-3_BT (**Supplemental Fig. S3A**). Because ruminant LINEs are generally longer than ancestral LINEs (**Supplemental Fig. S1C**), we hypothesized that long insertions are more strongly selected against within gene body regions. To test this, we classified ruminant LINEs into long and short groups (**Supplemental Fig. S3B**). Unlike L1_BT and L1-3_BT, long BovB insertions exhibited significant longer distance from the nearby genes (Wilcoxon test, p-value < 0.0001) than the short BovB insertions (**Supplemental Fig. S3C**), suggesting stronger selective constraints against long BovB insertions within gene-rich regions.

Notably, in contrast to L1_BT and L1-3_BT, no full-length BovB insertions were identified in the bovine genome. To investigate the mechanism underlying BovB truncation, we analyzed the relationship between ruminant LINEs insertion length and their sequence divergence from the consensus, which is a proxy for insertion age. The results revealed that TE insertion length was negatively correlated to sequence divergence for both L1_BT and L1-3_BT (Pearson r= -0.21 and -0.2, p-value < 2.2e-16, respectively), while only very weak correlations (Pearson r = -0.093, p-value < 2.2e-16) were obtained for BovB elements (**Supplemental Fig. S3D**). Thus, truncation of BovB elements appear to be independent from their age, suggesting that it is an inherent characteristic of their transposition process rather than a selective removal of longer insertions during evolution, which is the prevalent force observed in mammalian L1 elements (Levin et al., 2025), including the bovine L1 elements. Collectively, these findings reveal that ruminant-specific TEs, particularly BovB, exhibit peculiar genomic distribution patterns that differ from that of mammalian L1s which likely reflect both lineage-specific transposition dynamics and selective constraints in the bovine genome.

### Delayed ZGA and disrupted developmental trajectories in SCNT embryos

Given the distinct genomic organization of bovine TEs, we next investigated global transcriptional dynamics during bovine preimplantation development to establish developmental contexts for subsequent TE expression analyses. We reanalyzed publicly available RNA-seq datasets from *in vivo* fertilized (IVV), *in vitro* fertilized (IVT), and somatic cell nuclear transfer (SCNT) embryos spanning major developmental stages from MII oocytes to expanded blastocysts (**Supplemental Table S4)**. Gene-level and TE family-level expression (**Supplemental Tables S5-7**) was quantified simultaneously using TEtranscripts (Jin et al., 2015). After variance stabilizing transform and batch effects correction, samples were clustered into pre-ZGA and post-ZGA groups based on genes and TE families’ expression profiles (**Supplemental Fig. S4A-D**).

Consistent with previous studies (Graf et al., 2014; Hu, B. et al., 2024; Jiang et al., 2014), major ZGA occurred between the 4-cell to 8-cell stages in IVV embryos, whereas it was delayed to the 8-cell to 16-cell stages in both IVT and SCNT embryos (**Supplemental Fig. S4E, Tables S8-10**). ZGA genes were defined as lowly transcribed (< 50 read counts) in oocytes (2-cell stage for SCNT), significantly upregulated in 4-to 16-cell stages, and downregulated from the morula onward. Using this criterion, we identified 300, 262, and 270 ZGA genes in IVV, IVT, and SCNT embryos, respectively (**Fig. 2A, Supplemental Table S11**), with only 5 ZGA genes shared among all three embryo types, including *PNRC1, ARGFX, DUXA, LOC538435,* and *ZIM3* (**Fig. 2B and Supplemental Fig. S5A-C**). Of these, *DUXA*, *ZIM3,* and *ARGFX* are known transcription factors with critical roles during mammalian ZGA (Deng et al., 2020; Guo et al., 2025; Shi et al., 2025; Yaşar et al., 2025).

**Fig. 2.**
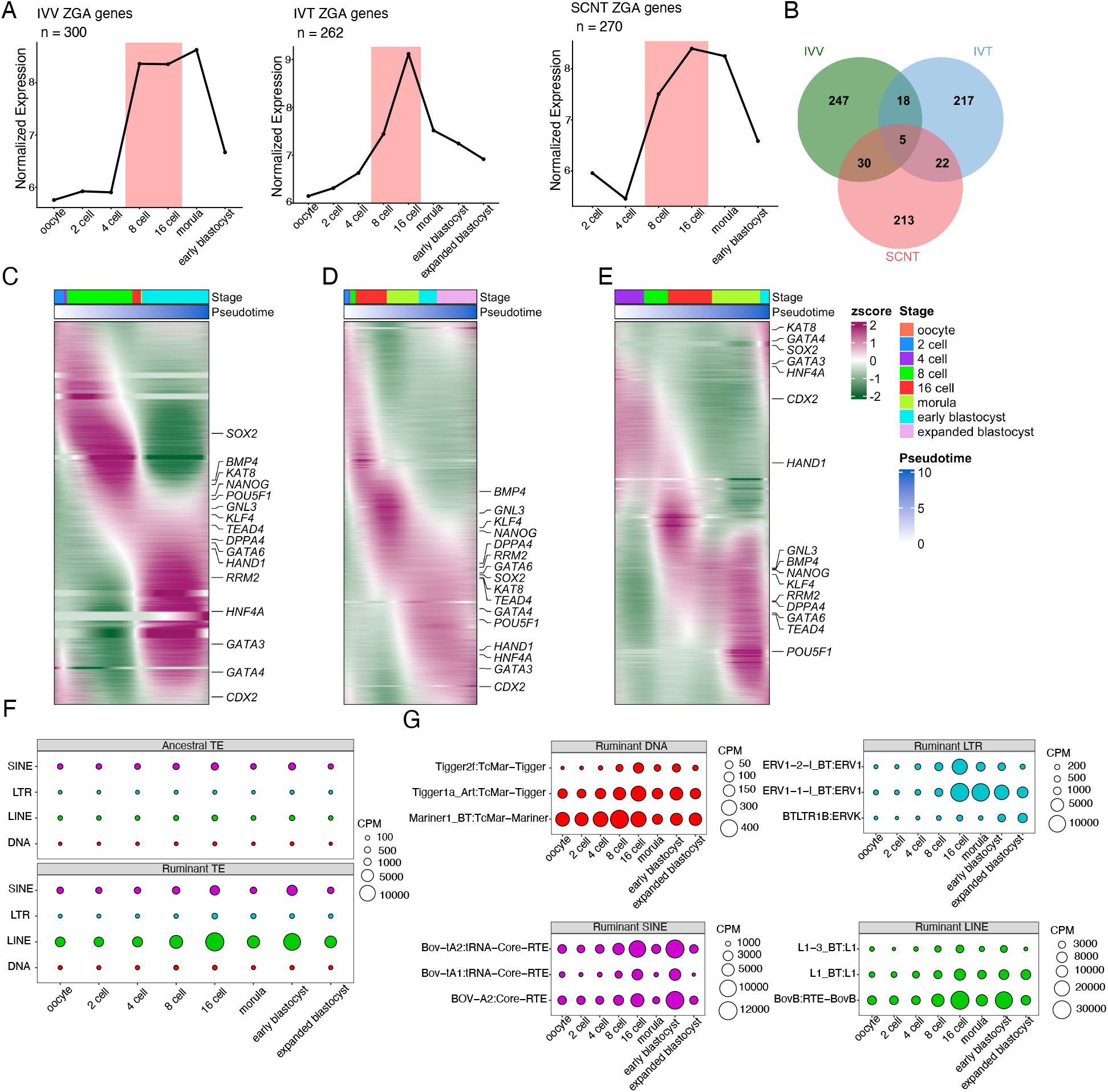
Gene expression dynamics and TE family expression. A: Mean expression level of ZGA genes in IVV, IVT and SCNT embryos. The vertical bar in line plot indicates the major ZGA stage in bovine embryos. B: Venn diagram indicates the overlapped ZGA genes across different embryo types. C: Gene expression heatmap ordered by pseudotime trajectory in IVV embryos. D: Gene expression heatmap ordered by pseudotime trajectory in IVT embryos. E: Gene expression heatmap ordered by pseudotime trajectory in SCNT embryos. Selected pluripotency genes and lineage markers are highlighted in heatmaps. F: Expression pattern of ancestral and ruminant TE family across preimplantation development. G: Expression pattern of top 3 TE families within each ruminant TE class.

To further characterize developmental progression, we reconstructed developmental psudotime trajectories using protein-coding gene expression. IVV and IVT embryos displayed continuous developmental progression from oocyte to blastocyst, whereas SCNT embryos exhibited disrupted transcriptional trajectories (**Fig. 2C-E and Supplemental Fig. S6A**). In addition, several pluripotency genes, including*, GNL3, HAND1, HNF4A, KAT8, SOX2,* and *POU5F1*, showed delayed activation in SCNT embryos (**Fig. 2E and Supplemental Fig. S6B-D**). In contrast, lineage genes (*CDX2*, *GATA3, GATA4,* and *HNF4A*) showed abnormal activation before ZGA (**Fig. 2E and Supplemental Fig. S6B-D**), consistent with incomplete developmental reprogramming in SCNT embryos. Together, these findings establish distinct transcriptional and developmental states across embryo types and provide a framework for investigating TE activation dynamics during bovine ZGA and its possible dysregulation in IVT and SCNT embryos.

### Ruminant-specific ERVs are coordinately activated with bovine ZGA

We next examined the extent to which TE expression coordinated with host genome gene expression during preimplantation development. Overall, we observed that ruminant-specific LINEs, SINEs, and LTRs exhibited substantially higher expression than older TEs across embryo development (**Fig. 2F**), including prominent families such as BovB, L1_BT, Bov-tA2, BOV-A2, and ERV1-1-I_BT (**Fig. 2G**). To identify coordinated TE-gene regulatory programs, we performed weighted gene co-expression network analysis (WGCNA) (Langfelder and Horvath, 2008) and identified 17, 14, and 12 co-expression modules in IVV, IVT, and SCNT embryos, respectively (**Supplemental Fig. S7**). Using these modules, we identified 32, 40, and 11 ZGA-associated TE families in IVV, IVT, and SCNT embryos, respectively (**Supplemental Fig. S8**, **Table S12**). All ZGA-associated TE families peaked at ZGA, followed by significant downregulation at later stages. Members of the ruminant-specific ERV1 and ERVL superfamilies were the predominant ZGA-specific TE families, with LTR45_BT and BtERV3B_LTR being commonly expressed at ZGA across all three embryo types (**Supplemental Fig. S8**). Together, these findings demonstrate that activation of ruminant-specific retrotransposons is tightly coordinated with ZGA-associated transcriptional programs during bovine early embryogenesis.

### Self-expressed retrotransposons undergo stage-specific activation

Building on the TE family and gene co-expression analysis, we next investigated stage-specific activation of individual TE loci during bovine preimplantation development. To distinguish autonomous TE transcription from passive co-transcription with nearby genes (Lanciano and Cristofari, 2020), TE loci were classified as either gene-dependent or self-expressed. Gene-dependent TE loci were defined as those overlapping with expressed gene bodies in the same orientation, while self-expressed TE loci were defined as those located within introns of non-expressed genes or in intergenic regions, therefore representing a set of TE loci more likely driven by their own regulatory elements (**Fig. 3A**). Across the three embryo types, we found that more than 60% of the total differentially expressed TE loci can be classified as self-expressed (**Fig. 3B and Supplemental Table S13**), indicating that these TEs possess promoters intrinsically activated at specific stages of early embryonic development. In contrast, differentially expressed gene-dependent TE loci, particularly LINEs, are predominantly enriched within 3’UTR and introns of actively transcribed genes (**Fig. 3C**), suggesting that these elements are co-expressed with developmentally regulated genes.

**Fig. 3.**
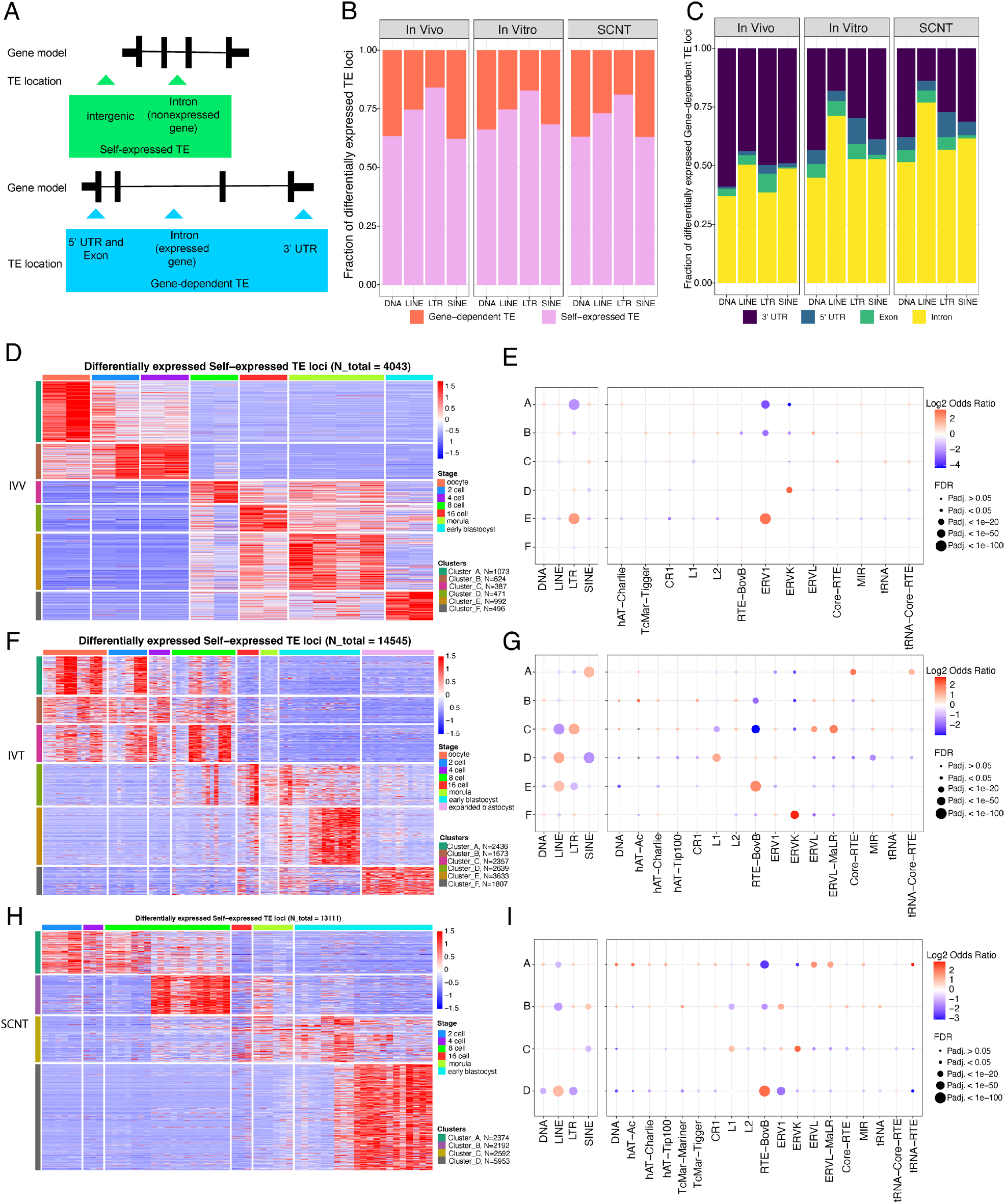
**TE loci classification and expression dynamics**. A: Definition of TE loci category. B: The proportions of TE loci category in different embryo types. C: The genomic location of active gene-dependent TE loci in different embryo types. D: Differentially expressed self-expressed TE loci of IVV embryo. E: TE class and family enrichment of IVV embryo. F: Differentially expressed self-expressed TE loci of IVT embryo. G: TE class and family enrichment of IVT embryo. H: Differentially expressed self-expressed TE loci of SCNT embryo. I: TE class and family enrichment of SCNT embryo. The cluster in enrichment plot corresponds to the clusters of differential expression heatmap.

Clustering analysis revealed highly dynamic and stage-specific activation patterns of self-expressed TE loci across embryo development (**Fig. 3D-I**). In IVV embryos, maternally deposited TE transcripts were progressively cleared after fertilization (**Fig. 3D**, clusters A & B), followed by robust activation of self-expressed TEs during the 4-to 8-cell transition (**Fig. 3D**, cluster C), corresponding to the major wave of ZGA. Distinct waves of TE activation were subsequently observed at 16-cell, morula, or blastocyst stages (**Fig. 3D**, clusters D-F). ERV elements, particularly the ERVK and ERV1 superfamilies, were activated at ZGA and beyond, predominantly enriched at the 16-cell and morula stages, respectively (**Fig. 3E**). In contrast, IVT embryos exhibited delayed activation of self-expressed TE loci, with major activation shifted towards the 8-to 16-cell stages (**Fig. 3F**, clusters C-D). SCNT embryos showed greater disruption, including weakened transcriptional activation of ZGA-associated self-expressed TEs (**Fig. 3H**) and abnormal dynamics of ERVL and ERVL-MaLR elements, which peaked at the 2-to 4-cell stages before ZGA but failed to persist later (**Fig. 3I**). In addition, IVT and SCNT embryos shared a progressive enrichment of LINE expression, including L1 and RTE-BovB elements, in post-ZGA development; a pattern not observed in IVV embryos (**Fig. 3E**, **Fig. 3G, and Fig. 3I**).

Overall, these findings demonstrated that self-expressed TE loci undergo highly coordinated and stage-specific activation during bovine embryogenesis, and that disruption of these TE activation programs is associated with delayed ZGA or abnormal genome reprogramming in IVT and SCNT embryos. These results further suggest that specific TE loci may function as regulatory elements contributing to transcriptional control during ZGA.

### DUXA-associated MLT1D activation during ZGA

To begin investigating the regulatory control of TEs activated at ZGA, we focused on the transcription factor *DUXA*, which out analysis identified as a ZGA marker across all embryo types (**Fig. 2B**). *DUXA* belongs to the double homeobox (DUX) gene family, whose homologs in mouse (*Dux*) and human (*DUX4)* directly activate a subset of TEs and genes at ZGA (De Iaco et al., 2017; Hendrickson et al., 2017; Whiddon et al., 2017). In cattle, *DUXA* is the only annotated DUX-family member previously proposed to regulate bovine ZGA (Leidenroth and Hewitt, 2010; Yaşar et al., 2025). Because epigenomic datasets were most extensively available for IVT embryos (Halstead et al., 2020; Ming et al., 2021; Zhou et al., 2023), subsequent analyses focused on this embryo type.

As expected, we observe that *DUXA* expression increased from the 4-cell stage and peaked at 8- and 16-cell stages, coinciding with major ZGA (**Fig. 4A**). To identify TE families potentially activated by DUXA, we performed TE loci enrichment analysis within the predicted DUXA binding regions of bovine genome. We found that MLT1D elements, which are LTRs from the ERVL-MaLR superfamily, were among the most significantly enriched TE families with the DUXA motif (**Supplemental Table S14**). Consistent with DUXA-mediated activation, MLT1D family expression was highly ZGA-specific (**Fig. 4B**). At the locus-specific level, we found that more than 89% of MLT1D copies are classified as self-expressed across oocytes and preimplantation development (**Fig. 4C**). Within these self-expressed MLT1D loci, only a small subset (∼1%, 163/13,910 loci) exhibited stage-wise differential expression (DE MLT1D) across embryonic stages (**Fig. 4D**, **Supplemental Table S15**). In contrast, non-DE MLT1D remained transcriptionally silent across early development (**Fig. 4D**). Interestingly, more than 70% of DE MLT1D loci were far away from genes in large intergenic regions (>30kb, **Fig. 4E**) and over 90% of them were candidate solo LTR insertions in the genome, suggesting potential roles as distal regulatory elements during bovine ZGA. Among the 163 DE MLT1D loci, 122 loci formed an active expression cluster during pre- and major-ZGA stages, including 82 MLT1D loci that significantly upregulated at 8 cell stage, whereas only 41 loci were activated post-ZGA (**Fig. 4F**).

**Fig. 4.**
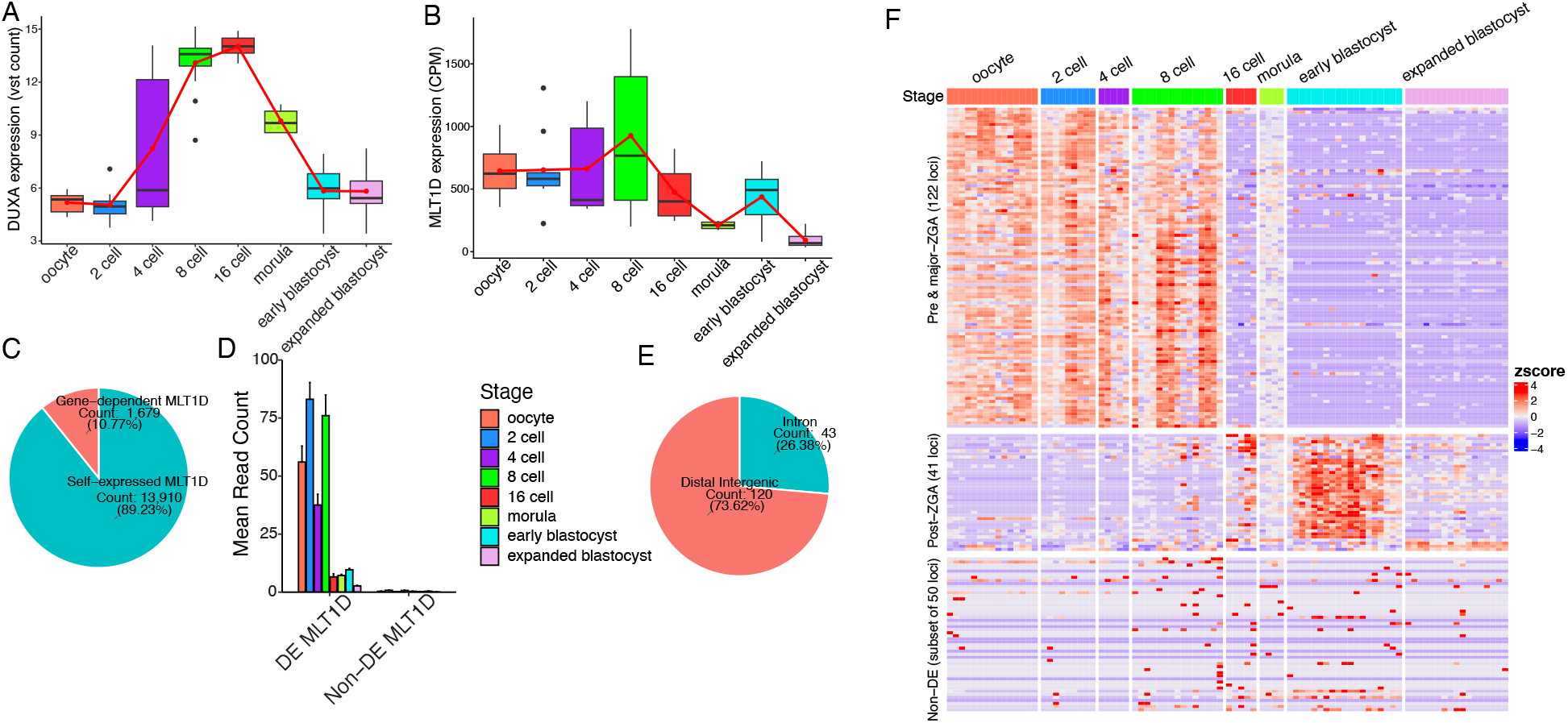
DUXA and MLT1D loci expression. A: DUXA expression in IVT embryo. B: MLT1D family expression in IVT embryo. C: Pie plot of gene-dependent and self-expressed MLT1D loci in IVT embryo. D. Expression patterns of differentially expressed MLT1D loci and non-differentially expressed MLT1D loci in IVT embryo. E: Pie chart shows the genomic distribution of differentially expressed self-expressed MLT1D loci in IVT embryo. F: Heatmap of MLT1D loci expression clusters.

To further validate autonomous transcriptional activation of these loci, we integrated 5’end RNA-seq data from early bovine embryos (Oomen et al., 2025). Strong transcription start site (TSS) signals were detected within the DE MLT1D loci, specifically during pre- and major-ZGA stages, whereas post-ZGA and non-DE MLT1D loci lacked detectable TSS activity (**Supplemental Fig. S9**). Together, these results suggest that a distinct subset of self-expressed MLT1D loci undergoes stage-specific activation during bovine ZGA. Specifically, these MLT1D elements are enriched for the DUXA-binding motif, located primarily in distal intergenic regions, and autonomously transcribed from their own promoters.

### ZGA-activated MLT1D loci acquire enhancer-like chromatin features

To infer the functional role of MLT1D elements at ZGA, we profiled the histone hallmark signals over all self-expressed MLT1D loci at 8 cell stage. Specifically, MLT1D loci exhibited high enrichment of both H3K27ac and H3K4me3 were defined as putative promoters, while MLT1D loci with high H3K27ac signals but lower or undetectable H3K4me3 signals were defined as putative enhancers (See Methods for details). Our results revealed that around 67% of the self-expressed MLT1D loci gained putative enhancer histone marks at ZGA (**Supplemental Table S16**). Especially, for the 82 MLT1D loci that showed significant expression upregulation at 8 cell stage, 58 of them were classified as putative enhancers according to the histone enrichment definition. Through further chromatin profiling of the 82 ZGA-expressed MLT1D loci, we confirmed that the activated MLT1D loci during bovine ZGA showed much higher enrichment of H3K27ac than the genome background, together with high chromatin accessibility and no signals of repressive marks (H3K27me3 and H3K9me3) at 8 cell stage (**Fig. 5A-G**).

**Fig. 5.**
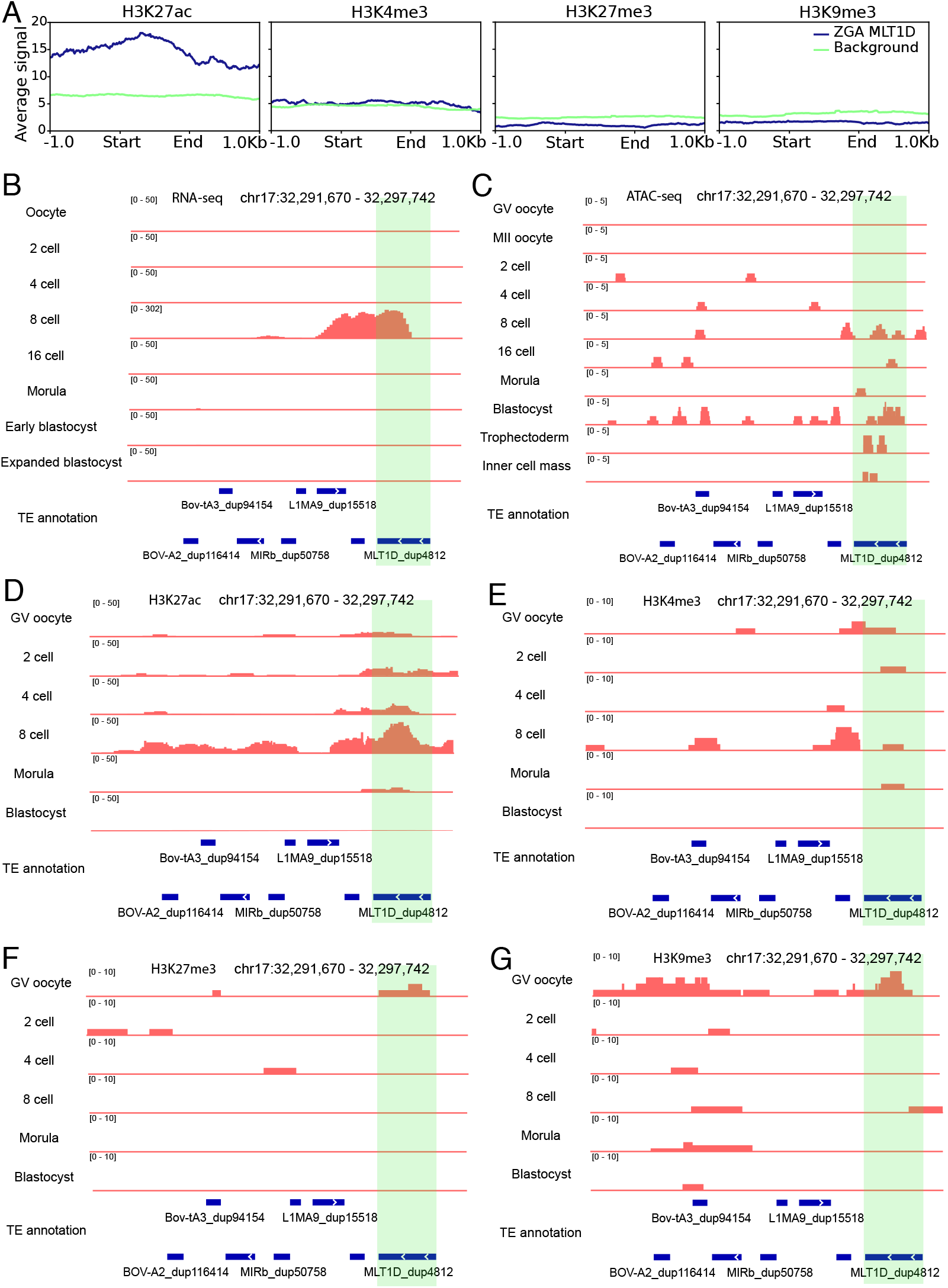
Epigenomic feature of ZGA MLT1D loci in IVT embryos. A: Histone enrichment over MLT1D loci specifically activated at 8 cell stage of ZGA. B: RNA-seq expression tracks of an 8 cell-specific MLT1D locus in IVT embryos. C: ATAC-seq tracks of an 8 cell-specific MLT1D locus in IVT embryos. D: H3K27ac tracks of an 8 cell-specific MLT1D locus in IVT embryos. E: H3K4me3 tracks of an 8 cell-specific MLT1D locus in IVT embryos. F: H3K27me3 tracks of an 8 cell-specific MLT1D locus in IVT embryos. G: H3K9me3 tracks of an 8 cell-specific MLT1D locus in IVT embryos.

Moreover, chromatin profiling of the MLT1D expression clusters in oocytes and preimplantation embryos revealed that the pre- and major-ZGA MLT1D cluster (n=122 loci) exhibited high H3K27ac and H3K4me3 enrichment in GV oocytes, followed by transient reduction during early cleavage stages and strong H3K27ac reactivation at the 8-cell stage (**Supplemental Fig. S10 A-B**). In contrast, H3K27me3 was accumulated over the pre- and major-ZGA MLT1D cluster in blastocyst stage (**Supplemental Fig. S10C**), whereas H3K9me3 was preferentially enriched at the post-ZGA MLT1D cluster (n=41 loci) in GV oocytes and pre-/major-ZGA embryos (**Supplemental Fig. S10D**), consistent with their repressive chromatin states. Moreover, the chromatin accessibility patterns of the MLT1D expression clusters closely aligned with the corresponding transcription activities across the embryonic stages (**Supplemental Fig. S11**). Together, our results support the cis-regulatory potential of activated MLT1D loci during bovine ZGA, which acquire H3K27ac-enriched active chromatin to function as enhancer-like elements.

### DUXA knockdown suppresses ZGA-Associated MLT1D Activation

To validate the functional relationship between DUXA and MLT1D loci, we first confirmed the presence of the DUXA motif within the MLT1D consensus sequence (**Fig. 6A**), suggesting that this motif is ancestral to MLT1D elements, i.e. it was present at the time of insertion of these elements in the bovine genome. We then performed de novo motif analysis over the pre- and major ZGA, the post-ZGA, and the non-DE MLT1D expression clusters. Interestingly, only motifs derived from the pre- and major ZGA MLT1D cluster showed a strong match with the DUXA motif (JASPAR ID MA0884.1, **Fig. 6A**). To further infer evolutionary origin of DUXA motif within MLT1D loci, we performed multi sequence alignment using the genomic sequences from pre-/major ZGA MLT1D loci and a random subset of 200 non-DE MLT1D loci. The phylogenetic analysis showed that pre-/major ZGA and non-DE MLT1D elements were interspersed across the tree rather than forming a single well-supported monophyletic clade (**Supplemental Fig. S12A**). Despite this phylogenetic dispersion, the DUXA motif was more frequently retained in ZGA-associated MLT1D elements and showed higher sequence conservation than in non-ZGA elements. Non-DE MLT1D copies more often exhibited partial loss of the homologous motif-containing region or sequence divergence within the motif (**Supplemental Fig. S12B-C**). Collectively, these results are more consistent with differential retention or loss of the DUXA motif across multiple MLT1D lineages than with acquisition of the motif by a single recently derived subfamily.

**Fig. 6.**
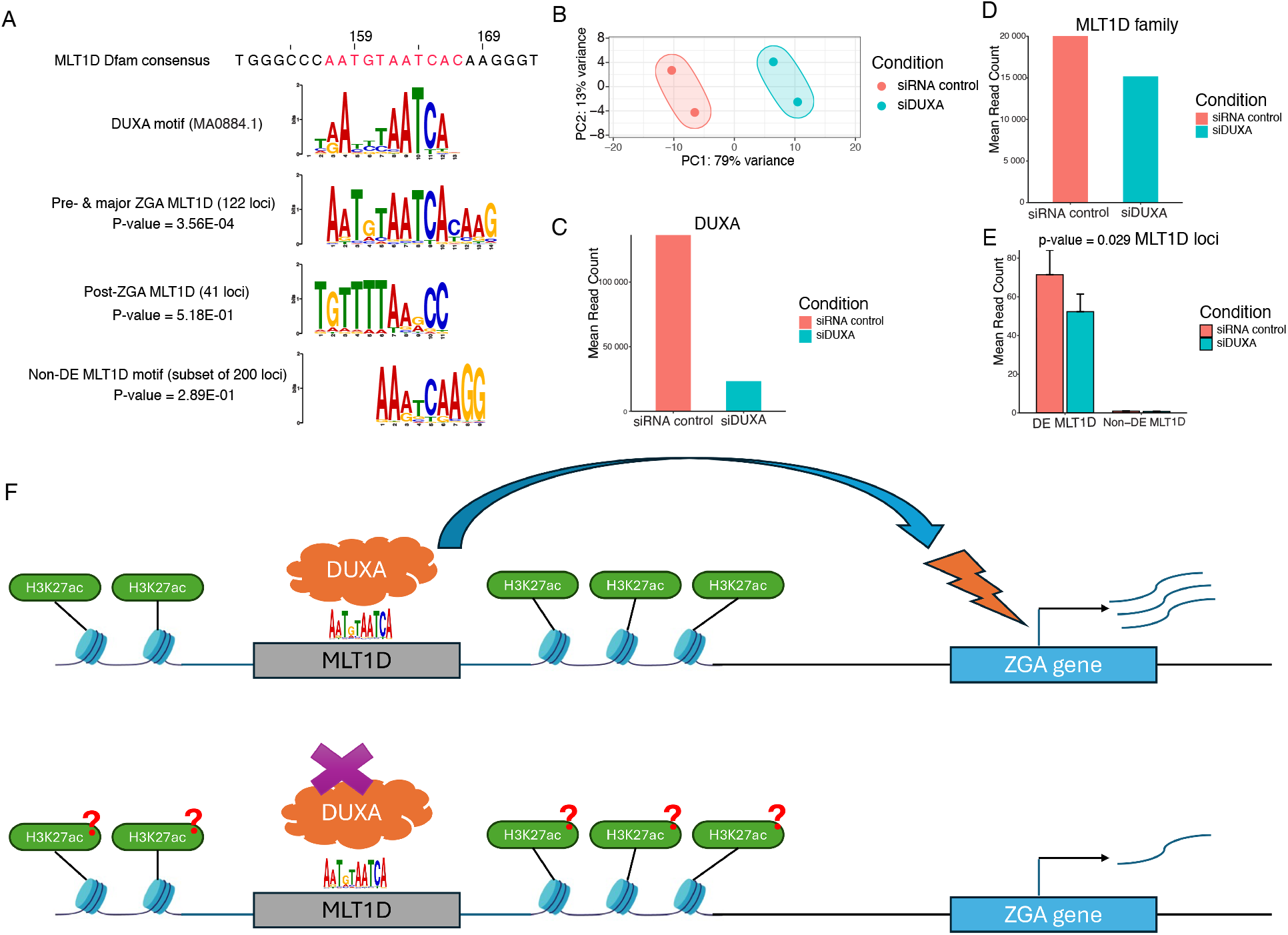
DUXA-MLT1D interaction and function validation. A: Top enriched DNA motif from different MLT1D expression clusters and DUXA motif overlapping analysis. B: PCA plot of DUXA knock down sample. C: DUXA expression under siRNA treatment. D: Expression changes of MLT1D family under DUXA siRNA treatment. E: Expression changes of differentially expressed MLT1D loci under DUXA siRNA treatment. F: Model for how DXUA and MLT1D function together to promote bovine ZGA.

To functionally assess DUXA-dependent regulation of MLT1D, we reanalyzed an RNA-seq dataset from *DUXA* knockdown experiments in 8-cell IVT embryos (Zhang et al., 2025). siRNA-mediated knockdown efficiently reduced *DUXA* expression (**Fig. 6B-C**) and resulted in 246 significantly downregulated genes, including several ZGA genes such as *DUXA, CD274, MAP3K8, MDFIC*, and *RCSD1* (**Supplemental Table S17-18**). At the TE family level, MLT1D family expression was significantly reduced following DUXA knockdown (**Fig. 6D**). Furthermore, expression of DE MLT1D individual loci was significantly decreased, whereas the expression of non-DE MLT1D loci remained unchanged (**Fig. 6E**). Together, these findings provide further evidence that DUXA directly regulates a distinct subset of ZGA-associated MLT1D loci during bovine embryogenesis.

## DISCUSSION

A growing body of literature suggest that TEs are not merely passive genomic passengers but active components of genome function. Notably, there is now ample evidence that specific groups of TEs can function as coordinated cis-regulatory elements orchestrating the complex gene regulatory networks required for embryogenesis (Deaville and Berrens, 2024; DiRusso and Clark, 2023; Fueyo et al., 2025; Guo et al., 2024). Recent advances in multi-omics and integrative genomic analysis have further revealed the pivotal roles of TE activities during embryogenesis in human and mouse (Fueyo et al., 2022; Li et al., 2026), yet their regulatory functions during early development in other mammals, including livestock such as cattle, remain poorly studied.

By combining genomic, transcriptomic, and epigenomic analysis, our study represents, to our knowledge, the most comprehensive characterization of bovine TEs during preimplantation embryogenesis. Previous studies primarily described stage-specific activation of several ERV families during preimplantation development (Bui et al., 2009; Khazaee et al., 2018; Oomen et al., 2025). Our work significantly expands on these studies by integrating the evolutionary history, genomic organization, and developmental activation dynamics of both ancestral and ruminant-specific TEs across multiple embryo types. Notably, we identified distinct genomic distribution and expression patterns for ruminant-specific TE families and uncovered stage-specific activation of a large pool of TE loci during bovine ZGA. Consistent with previous reports in mammals (Guo et al., 2024; Li et al., 2026), ERV/LTR families represented the predominant class of TEs activated during preimplantation development, even though these elements have distinct evolutionary origins (ERV1, ERVL, ERVK) and have infiltrated mammalian genomes independently. It suggests that ERVs possess intrinsic features, such as cis-regulatory sequences, that predispose them to be active in early mammalian embryos. Presumably, these features enabled them to transpose in the germline, but even prior to the specification of germ cells, and colonize successfully mammalian genomes.

By comparing IVV, IVT, and SCNT embryos, we further demonstrated that TE activation closely mirrors the timing of ZGA and become disrupted in SCNT embryos, suggesting the intriguing possibility that proper TE activation may be needed for successful developmental reprogramming of bovine embryos, as suggested by other studies in mouse and human models (Fueyo et al., 2025; Hendrickson et al., 2017; Xiang et al., 2026). Importantly, our study identified a distinct subset of ZGA-activated, self-expressed MLT1D loci enriched for the DUXA-binding motif. RNA-seq analysis of DUXA-depleted embryos revealed a significant reduction of MLT1D activation during ZGA, supporting a regulatory relationship between DUXA and MLT1D activation. These DUXA-regulated MLT1D elements are predominantly represented by solo LTRs located far away from genes in vast intergenic regions and they are marked by highly accessible chromatin and H3K27ac during ZGA, suggesting that they may act as distal enhancers for coordinating bovine gene expression at ZGA (**Fig. 6F)**. Moreover, we confirmed the potential of MLT1D for initiating chimeric transcripts by mapping transcription initiation signals within these MLT1D loci. Importantly, another ERVL element, MLT2A1, has recently been proved for producing chimeric transcripts to regulate human ZGA (Xiang et al., 2026), which indicating an intriguing convergence between human and bovine. Together, these findings support a hypothesis in which specific ERV loci function as regulatory elements during bovine ZGA and highlight the intertwinement of TEs, transcription factors, and chromatin remodeling during early embryonic genome activation.

### Ruminant-specific TEs reshape the bovine genomic landscape

TEs account for around 47% of the bovine genome, with LINEs representing the most abundant class, on par with the TE content of other mammals (Osmanski et al., 2023; Wang et al., 2025; Zhao et al., 2023). Nonetheless, a substantial fraction (∼ 25%) of TEs in the bovine genome are ruminant-specific: they inserted after the split of ruminants from other mammalian lineages such as the rodent and primate lineages, approximately 50 My ago. In fact, some TE families show very low intra-family sequence divergence, indicative of very recent transposition activity. Consistent with recent reports of ongoing mobilization in the cattle germline (Tang et al., 2024), we identified ERVK as the youngest TE superfamily in the bovine genome. Several young LINEs and SINEs families were also detected, reflecting the continuous activity of LINEs in almost all mammalian species and the known dependence of SINEs retrotransposition on the LINE-encoded machinery (Kramerov and Vassetzky, 2011).

At the genome-wide level, bovine TE chromosomal distributions share conserved features with human and mouse genomes, including a positive correlation between the chromosomal density of SINEs and DNA transposons and a negative correlation between LINEs and SINEs (Chinwalla et al., 2002; Lander et al., 2001). However, when separating ancestral and ruminant TEs, we observed previously unrecognized genome features more specific to the bovine genome. For instance, BovB elements constituted a much higher proportion for ruminant-specific LINEs than L1 elements, probably due to their massive genomic amplification in the bovine following horizontal introduction in the ruminant lineage (Ivancevic et al., 2018; Kordis and Gubensek, 1998, 1999). Particularly, we found ruminant-specific LINEs were predominantly located in intergenic regions, whereas ancestral LINEs retained localization across gene body regions, suggesting potential purifying selection against younger LINE insertions within genes in bovine as reported for LINE-1 elements in human and mouse (Boissinot et al., 2001; Graham and Boissinot, 2006). In addition, canonical correlations among TE classes observed with ancestral mammalian TEs were partially lost when ruminant-specific TEs were considered, such as the positive correlation between DNA transposon and LINEs and SINEs, indicating substantial lineage-specific reorganization of TE genomic architecture during ruminant evolution.

### Distinct ZGA regulatory programs and TE activation dynamics across embryo types

Successful preimplantation development requires precise temporal control of gene expression and coordinated regulation of noncoding elements during ZGA (Tian et al., 2020). By systematically comparing IVV, IVT, and SCNT embryos using unified criteria, our study refined bovine ZGA-associated genes and identified shared and embryo-type-specific regulatory factors. Among the shared ZGA regulators, DUXA, ZIM3, and ARGFX were consistently activated across the three embryo types. DUXA belongs to the conserved DUX-family pioneer transcription factors that promote ZGA and pluripotency establishment in multiple mammals (De Iaco et al., 2017), including DUX4 in humans (Vuoristo et al., 2022), Dux in mice (De Iaco et al., 2020), and DUXA in porcine SCNT embryos (Shi et al., 2025). Moreover, a comparative study across pig, cow, rabbit, mouse, and rhesus revealed that DUX family members were activated during ZGA and shared common binding motifs enriched at ZGA genes in all species examined (Oomen et al., 2025). Consistent with these studies, our results showed DUXA expression peaked at major ZGA at 8 and 16 cell stages, supporting a conserved role in mammalian ZGA. In addition to DUXA, our analyses identified ZIM3 and ARGFX as novel candidate bovine ZGA regulators. ZIM3 is an imprinted gene in bovine (Kim et al., 2007) and has been implicated as a potential ZGA regulator in humans (Li, J. et al., 2025) and goats (Deng et al., 2020). ARGFX was also shown to regulate cleavage-stage development and ZGA-associated transcription in human embryos (Guo et al., 2025).

We further identified embryo-type-specific candidate regulators, including OTX1 in IVV embryos, multiple ZNF-family transcription factors in IVT embryos, and KLF17 in SCNT embryos. OTX1 is critical for ZGA in Xenopus early embryos (Paraiso et al., 2019), while ZNF family members, especially the KRAB domain-containing ZFPs, also play an important role in bovine embryo development (Zhang et al., 2022). KLF17 was also known to recruit RNA Pol II to initiate ZGA genes in mouse embryos (Hu, Y. et al., 2024) and found to be a minor ZGA gene in bovine early development (Zhang et al., 2025). These findings suggest that altered usages of ZGA regulators networks may contribute to the delayed genome activation and reduced developmental competence under IVT and SCNT compared to IVV embryos, and further experimental studies are required to validate this hypothesis.

Beyond ZGA regulators, our study also revealed coordinated activation of specific TE families during bovine ZGA. Previous studies in mouse and human demonstrated that ERVL-associated retrotransposons, including MERVL and HERVL, respectively, are transiently activated during ZGA and a subset of these elements likely function as regulatory elements promoting early embryonic transcription (Göke et al., 2015; Liu et al., 2019; Macfarlan et al., 2012; Peaston et al., 2004; Pontis et al., 2019; Svoboda et al., 2004; Xiang et al., 2026). Consistent with these conserved mammalian patterns, we identified ERV1, ERVL, and L1 families as the predominant TE groups activated at bovine ZGA. Importantly, our study further demonstrated embryo-type-specific TE activation dynamics and widespread activation of self-expressed TE loci during ZGA. Through comparative analysis across three embryo types, we observed greater abundance of LTR and LINE activation in IVV and IVT embryos than SCNT embryos, including ERV1-2-LTR_BT, ERV1-1-I_BT, and X15_LINE elements. Together, these findings support a model in which coordinated activation of both ZGA transcription factors and TE regulatory programs is required for successful mammalian embryonic reprogramming. Disruption of these coordinated programs in SCNT embryos may therefore represent a major barrier to normal developmental progression.

### Stage-specific activation of self-expressed TE loci in bovine embryogenesis

Although quantifying TE expression from short-read RNA-seq remains challenging because of repetitive sequences and overlap with host transcripts (Kapusta et al., 2013; Lanciano and Cristofari, 2020), our locus-level analyses enabled separation of TEs into gene-dependent and self-expressed categories. More than half of the differentially expressed loci were self-expressed, suggesting widespread autonomous TE transcription during bovine embryogenesis. Distinct TE classes also exhibited highly stage-specific activation patterns. For example, self-expressed SINEs and DNA transposons were abundant in oocytes and 2-cell stage but depleted following ZGA, whereas LTRs became more active at ZGA, consistent with observations in other mammals (Li, B. et al., 2025; Yang et al., 2024).

Importantly, our comparative analyses further revealed significant embryo-type-specific differences in TE activation dynamics. In IVV embryos, ERVK activation coincided with the major ZGA, whereas in IVT and SCNT embryos, it was delayed until post-ZGA stages. Moreover, ERVL and ERVL-MaLR superfamilies, which are normally activated during ZGA, showed premature activation at the 2-cell stage but failed to maintain activation during ZGA in SCNT embryos. Because transient ERV transcription is recognized as a critical component in successful ZGA (Fu et al., 2019; Sakashita et al., 2023), these abnormal TE activation patterns may reflect incomplete epigenomic reprogramming and contribute to the reduced developmental potential in bovine SCNT embryos (Grow et al., 2023).

### MLT1D elements as candidate cis-regulatory elements during bovine ZGA

Among the activated TE families, MLT1D emerged as a strong candidate regulator associated with bovine ZGA. In our study, MLT1D loci were significantly enriched for the DUXA binding motif. Differential analysis identified 163 stage-specific DE MLT1D loci, near three-quarters of which were highly expressed in pre-and major-ZGA stages. Consistent with autonomous transcriptional activation, strong transcription initiation signals were captured within activated MLT1D loci during early development, which also support previous reports that MLT1D elements can initiate chimeric RNAs in bovine preimplantation development (Oomen et al., 2025). Moreover, a recent publication in human embryonic genome activation reveals that MLT2A1, a member of ERVL elements, could form chimeric transcripts to interact with ZGA genes and recruit co-factors for facilitating transcription (Xiang et al., 2026). As another member of ERVL elements, whether MLT1D play a ruminant-specific role in ZGA or following similar mechanism as human can be further explored using long-read sequencing.

Epigenomic analyses further illustrated a regulatory role for activated MLT1D loci. Although the enrichment signal of H3K27ac was twofold higher than H3K4me3 signal in GV oocytes over pre-and major-ZGA MLT1D loci, both histone marks had higher enrichment than random genome background, suggesting weak promoter features of MLT1D loci at this stage. During major ZGA at 8 cell stage, H3K4me3 was largely removed while H3K27ac was broadly deposited over pre- and major-ZGA MLT1D loci. Combined with their distal intergenic localization and dynamic chromatin accessibility, these findings suggest that a subset of MLT1D element act as enhancer-like elements at bovine ZGA. However, more studies are needed to confirm if MLT1D elements play different cis-regulatory roles across embryonic development.

In addition, *de novo* motif analysis identified DUXA as the top-enriched TF specifically in pre- and major-ZGA loci instead of post-ZGA loci, which may be due to differential motif preservation in certain MLT1D lineages according to the phylogeny analysis. Functional analysis of DUXA knockdown embryos further demonstrated impaired activation of MLT1D loci and ZGA genes during bovine ZGA. DUXA has been shown as one of the master regulators that facilitate bovine ZGA in previous studies (Leidenroth and Hewitt, 2010; Yaşar et al., 2025), but little info has been reported for the biological role of MLT1D in bovine embryogenesis. Collectively, our findings provided insights that MLT1D might interact with DUXA to promote transcriptional activation during bovine ZGA, but further functional experiments are needed to validate its direct impact in early development.

## CONCLUSION

This study provides the most comprehensive characterization to date of TE regulation during bovine embryogenesis by integrating genomic, transcriptomic, and epigenomic analyses across IVV, IVT, and SCNT embryos. Our findings reveal the molecular impacts of different embryo culture conditions and highlight the key factors contributing to zygotic genome activation. More broadly, our work establishes a foundational framework to identify TE-associated biomarkers and regulatory pathways linked to embryo quality, developmental competence, and epigenetic reprogramming to improve livestock reproduction.

## METHODS

### Transposable element annotation and age estimation

Bovine reference genome (ARS-UCD 2.0) was downloaded from NCBI database and TE mapping was conducted using the rmblastn engine (v 2.14.0+) of RepeatMasker software (v 4.1.5) (Smit AFA, 2013-2015), together with the Dfam (Storer et al., 2021) consensus and Repbase-20181026 libraries (Bao et al., 2015). The following parameters were set: -a, -s, -nolow, -gccalc, -gff, -cutoff 200, -no_is -species cow. Based on the RepeatMasker output, the Kimura CpG-corrected percentage-divergence from the consensus sequence was computed using ParseRM (Kapusta et al., 2017). The TE copy number estimates were obtained using the Perl script one_code_to_find_them_all.pl (Bailly-Bechet et al., 2014), including the reconstruction of fragmented repeats and full-length LTR elements. For defining the ancestral TE and ruminant TE, the evolutionary age of TE families was extracted from the Dfam annotation (Storer et al., 2021) using the famdb tools from github (https://github.com/Dfam-consortium/FamDB). TE families detected in the ancestors of ruminant lineage were determined as ancestral TE, while the rest were determined as ruminant TE.

To estimate the insertion age of each TE family, defragmented sequences were extracted from the genome, while a minimum sequence length of 100 was used for further analysis (Chang et al., 2022). Multiple sequence alignment was performed using MAFFT (v7.520) (Kuraku et al., 2013) with the --auto flag equal to true, and the alignments were trimmed using trimAl (v 1.4) (Capella-Gutiérrez et al., 2009) with the -gt parameter equal to 0.01. TEs with less than 10 sequences were ignored, and alignments of TEs with a large copy number were subjected to a random subset of 1250 sequences for improving computation efficiency. Then, approximate maximum-likelihood phylogenetic trees were constructed using FastTree (v 2.1.11) (Price et al., 2010) with a generalized time-reversible model, and branch lengths were rescaled to optimize Gamma20 likelihoods. The age of individual TE insertion was obtained as the branch length from the leaf to the most recent ancestor (terminal branch length), and the age of a TE family was determined as the median of the terminal branch lengths. For evaluating the TE age estimation method, branch length-based methods were compared with the TE age calculated by RepeatMasker-derived divergence from consensus sequence.

### Genomic distribution analysis

To define the genomic distribution of TEs, we divided the bovine into non-overlapping 2-Mbp windows and calculated the genomic density for specified features in each window, including genes, satellite markers (BTSAT), GC nucleotide content, LTRs, LINEs, SINEs and DNA transposons. The density results were visualized using Circos (Krzywinski et al., 2009), and the correlations among the distributions of genes, satellite and different TE classes were computed with Spearman’s rank correlation on the windows, ordered by coverage density.

To assess if TEs were associated with functional genomic regions, we focused on TEs that overlapped with intergenic regions, gene body, exons, introns, and promoter regions (1kb ± TSS). Specifically, we calculated the distance between a TE locus and the targeted genomic regions, and a TE family would be defined as “preferentially located” in the target regions if the median distance between the target region and TE copies from that family was equal to zero. We then calculated the observed fraction of preferentially enriched TE families within each TE class. Next, we computed the expected fraction of preferentially enriched TE families in target regions through random shuffling of TE labels across the genome, which removed the differences between families but maintained the original TE distributions. We summarized the expected fractions of preferentially enriched TE families through 1,000 permutations for each TE class, and the significance of the discrepancy between observed and expected fractions was determined with binomial tests.

For evaluating TE distributions at different level, the same analysis process was applied to overall TEs, ancestral TEs and ruminant TEs separately. For splitting the long and short insertions in ruminant LINE, 1500 bp, 4200 bp, 4250 bp were used as length thresholds for BovB, L1_BT, and L1-3_BT, respectively. The distance between TE insertions and the nearest genes was calculated using bedtools closest function with -d -s parameter, and the difference of median distance between long and short insertions was calculated using Wilcoxon rank-sum test. For checking the correlation between insertion length and sequence divergence for each ruminant LINE family, the sequence divergence and sequence length of TE insertion were obtained from RepeatMasker annotation output to calculate the Pearson correlation coefficient.

### RNA-seq data pre-processing

In this study, a total of 163 RNA-seq samples were selected from NCBI GEO database, including 16 IVV samples, 89 IVT samples and 58 SCNT samples. The low-quality reads (percentage of Q20 bases < 60%) and sequencing adapters were removed from raw data by fastp (Chen et al., 2018). Clean reads were aligned to bovine reference genome using STAR (Dobin et al., 2013) with the specified parameters: --chimSegmentMin 10 --winAnchorMultimapNmax 200 --outFilterMultimapNmax 100. The output bam files were sorted and indexed using samtools (Li et al., 2009). Then the TEcount function of TEtranscripts (Jin et al., 2015) was applied to get gene counts and family-level TE counts with the following parameters: --stranded no –sortByPos -- mode multi. For exploring the sample relationships, VST transformation and SVA correction were performed with the raw gene counts, and Pearson correlation coefficients were computed across samples, followed by visualization using Complexheatmap (Gu et al., 2016).

### Differential gene expression and pseudotime analysis

To capture the gene expression dynamics in early development, differential gene expression was performed using DESeq2 (Love et al., 2014) with pair-wise comparisons between adjacent stages. Lowly expressed genes were removed and SVA batch correction was included in the experimental design to account for potential confounders. The DEGs were obtained with the specified threshold: padj < 0.05 and |log2 fold change (FC)| >1. Based on the DEGs across embryonic stages, ZGA genes were defined in each embryo type with the following criteria: low expression (read counts < 50) in oocytes (2 cell for SCNT), significant upregulation at 4 cell, 8 cell or 16 cell stages, significant downregulation in morula or early blastocysts, and no signs of upregulation in expanded blastocysts.

For pseudotime analysis, the average gene expression at each developmental stage was summarized from the VST-SVA corrected expression profile, and pseudotime analysis was performed using bulkPseudotime R package (Zhang, 2024). Briefly, principal component analysis (PCA) was performed to determine the Euclidean distance between the time points, which would be rescaled to a new pseudotime path between 0 ∼ 10. Then gene expression was imputed and ordered along the pseudotime trajectory, followed by visualization using Complexheatmap (Gu et al., 2016). Additionally, a total of 16 pluripotent genes and lineage specification markers were highlighted on the pseudotime heatmap and their expression patterns were further visualized with pseudotime line plots.

### Gene-TE co-expression analysis

To explore the relationship between genes and TEs in embryogenesis, co-expression analysis that covering both gene expression and TE family-level expression was conducted using WGCNA (Langfelder and Horvath, 2008). Nonexpressed genes and TE families were removed in the analysis, followed by VST-SVA correction for addressing potential batch effect when combining samples from different sources. We followed the official protocol for automatic network construction (Langfelder and Horvath, 2008), and soft-thresholding powers chosen for IVV, IVT and SCNT embryos were 9, 12, 12, respectively. We treated embryonic stages as the targeted traits and computed their Pearson correlations to the module eigengenes to screen for significant correlated modules with each stage. For quantifying associations of individual genes with the targeted traits, gene significance was defined as the correlation between the gene and the traits. Furthermore, we focused on the co-expression modules that exhibited significantly positive correlations (p < 0.05) with the bovine ZGA process, including 4 cell, 8 cell and 16 cell stages. Within these selected modules, TE families that were significantly (p < 0.05) correlated with ZGA stages were summarized and visualized using ComplexHeatmap.

### TE loci-level expression and classification

The locus-specific quantification of TE expression was achieved using the telescope assign function of Telescope (Bendall et al., 2019), together with the following parameters: --theta_prior 200000 --max_iter 200 --updated_sam. Counts from TE fragments reconstructed by one_code_to_find_them_all.pl were merged. To differentiate activate TE transcription from passive gene co-expression, we applied a similar method as previous studies (Chang et al., 2022) to classify the TE loci into gene-dependent and self-expressed categories. Briefly, the relative location of TE loci to protein-coding genes was annotated to different genomic features using ChIPseeker (Yu et al., 2015), including 5’ UTR, 3’ UTR, exon, promoter, intron, downstream and intergenic. Gene expression information was also included in the analysis, and genes with >=10 normalized counts in at least 1 sample will be defined as expressed genes, otherwise defined as nonexpressed genes. If TEs were overlapped with exons, 5’ and 3’ UTRs or the intron of expressed genes, they would be labeled as gene-dependent TEs. Conversely, TEs associated with intergenic regions or intron of nonexpressed genes were labeled as self-expressed TEs. For TE fragments reconstructed as components of the same TE through one_code_to_find_them_all.pl (Bailly-Bechet et al., 2014), they would be labeled in the same category based on the following hierarchy: exon > 3’ UTR > 5’ UTR > intron of expressed gene > intron of nonexpressed gene > intergenic.

### Differential TE loci expression analysis and enrichment analysis

For each embryo type, differential TE loci expression was conducted using DEseq2 (Love et al., 2014). The lowly expressed TE loci were removed in the analysis, which required >10 read counts at each stage. Due to the dominant proportion of gene-related read counts in embryonic transcriptome, TE loci counts were merged with gene counts when performing DESeq normalization, which could provide a better dispersion estimation. To capture TE loci with stage-specific expression profile, pairwise comparisons were applied between any developmental stages. Then Benjamini-Hochberg correction was conducted for the combined results from all pairwise comparisons using p.adjust function in R. Finally, only significant differentially expressed self-expressed TE loci were be considered in further analysis, which were selected based on adjusted P-value < 0.01 in any pairwise comparisons. For visualizing the TE expression pattern, Z-score standardization of the differentially expressed TE loci was presented in heatmaps, and kmeans clustering was performed with k = 6, 6, 4 in IVV, IVT and SCNT embryos, respectively. According to the TE expression clusters, fisher exact test was applied to check the enrichment of TE classes and families for each TE loci cluster, together with multiple testing to calculate the adjusted P-values.

### ATAC-seq analysis

ATAC-seq data in bovine preimplantation embryos were downloaded from NCBI GEO database (GSE143658 and GSE145040), including GV oocytes, MII oocytes, 2 cell, 4 cell, 8 cell, 16 cell, morula and blastocysts. Additionally, ATAC-seq data of trophectoderm and inner cell mass in bovine blastocysts were downloaded from NCBI with the accession number GSE193640. Briefly, sequencing adapter and low quality reads (q < 20) were removed from raw data using Trim_Galore (Felix, 2021), together with a minimum sequence length of 10bp. Clean reads were mapped to bovine reference genome (ARS-UCD 2.0) using BWA (Li et al., 2009) aln (-q 15) and sampe, followed by PCR duplicates removal with Picard (Institute, 2018) and low-quality (q < 15) alignments removal with SAMtools. The nucleosome-free region bam files were then generated using SAMtools with sequence length <= 100bp. For calculating the genome-wide ATAC-seq signals with deepTools (Ramírez et al., 2016), firstly all samples were analyzed together to calculate the scale factors using multiBamSummary with the 10bp bin, then the bigwig files were obtained using bamCoverage with the following setting: --scaleFactor –skipNonCoveredRigions --extendReads -bs 1. In terms of the biological replicates per stage, we merged the bigwig files using bigWigMerge and bedGraphToBigWig functions from UCSC_tools (Kent et al., 2010). Next, the summary statistics were calculated using computeMatrix with the following parameters: scale-regions -m 1000 -a 1000 -b 1000 -missingDataAsZero, and the ATAC-seq signal over the differentially expressed TE loci was visualized using plotProfile. For highlighting the changes in chromatin accessibility, we also plotted the ATAC-seq signal over the background region, which were corresponded to 1000 random regions within bovine genome with a length of 500bp.

### CUT&Tag analysis

CUT&Tag data of two active histone marks (H3K27ac and H3K4me3) and two repressive histone marks (H3K27me3 and H3K9me3) in bovine early embryos were obtained from NCBI GEO database (GSE193640), ranging from GV oocytes to blastocysts, as well as the associated trophectoderm and inner cell mass. Quality control of raw data was performed using Trim Galore with the following parameters: -q 20 --length 10 -stringency 1 --gzip. Clean reads were aligned to bovine reference genome using the mem module of BWA. PCR duplicates were removed using PicardTools (Institute, 2018) and low-quality mapping (q < 15) were eliminated with SAMtools. Then similar steps as ATAC-seq data were conducted for visualization using deepTools (Ramírez et al., 2016).

### 5’end RNA-seq analysis

5’end RNA-seq data in bovine preimplantation embryos were downloaded from NCBI GEO database (GSE225056), ranging from oocytes to 16 cell stage. Low quality reads and sequencing adapters were removed using trimmomatic (Bolger et al., 2014). Clean reads were mapped to bovine reference genome using STAR and unique mapping was selected using samtools. Only Illumina-format read2 was used for downstream analysis as this contains the true 5’ transcript information. rRNA reads were removed based on repeatmasker annotation, together with bedtools (Quinlan and Hall, 2010) and picard. The enrichment of 5’end transcripts were visualized using bedgraphToBigWig (Kent et al., 2010) and deepTools (Ramírez et al., 2016).

### DUXA and TE interaction analysis

The DUXA motif was obtained from JASPAR database (Rauluseviciute et al., 2024) and the potential DUXA binding regions in bovine genome were screened using FIMO (Grant et al., 2011). We explored the enrichment of TE family within the DUXA binding regions using TEanalysis suite (Kapusta et al., 2017). Briefly, we performed a binomial test to check if the number of overlaps between DUXA binding regions and TE families was significantly higher than expectations, which were derived by randomly shuffling TE positions on the same chromosome. Since MLT1D TE loci were significantly associated with DUXA, we plotted the expression pattern of MLT1D loci using Complexheatmap (Gu et al., 2016) and performed kmeans clustering to define pre- and major-ZGA cluster, post-ZGA cluster and non-DE cluster. The ATAC-seq signal and histone modifications over an 8 cell-specific MLT1D locus were visualized IGV genome browser (Robinson et al., 2011). The enrichment of ATAC-seq, histone and 5’end RNA-seq over MLT1D loci across embryonic stages were visualized using deepTools (Ramírez et al., 2016). To define the putative enhancers at 8 cell stage, we calculated the enrichment signal of H3K27ac and H3K4me3 for each self-expressed MLT1D locus using multiBigwigSummary function from deeptools. Then the putative enhancers were defined following criteria: only H3K27ac enrichment without H3K4me3 enrichment, or H3K27ac enrichment is 1.5 folds higher than H3K4me3 enrichment.

### Sequence analysis of MLT1D and DUXA function evaluation

For predicting the regulator of each MLT1D expression cluster, the fasta sequences of the MLT1D clusters were retrieved using bedtools getfasta function with -name -s parameters. Then the fasta sequences were used as inputs to XSTREME (Grant and Bailey, 2021) for obtaining the top enriched DNA motifs and comparing with DUXA motif (MA0884.1) from JASPAR database. To balance the input size for DNA motif prediction, we randomly selected 200 MLT1D loci from the non-DE MLT1D cluster for the motif analysis. For phylogeny analysis, genomic sequences from the pre- and major ZGA MLT1D loci and random non-DE MLT1D loci were aligned together with MLT1D consensus sequence from Dfam database using MAFFT (Kuraku et al., 2013). Then the large gaps in the alignment were removed using trimAl (Capella-Gutiérrez et al., 2009) with the option -gt 0.01. Finally, the maximum-likelihood phylogenetic tree was generated using FastTree (Price et al., 2010) with the parameters -nt -gtr. For comparing the DUXA motif coverage and identity between different MLT1D groups, the significance was computed using Wilcoxon test. To evaluate the functional relationships between DUXA and MLT1D, we reanalyzed a public dataset that performed DUXA knock down at 8 cell stages of bovine embryos. The differential gene expression and TE loci expression analysis were conducted using similar steps as bulk RNA-seq processing across different embryo types. We summarized the expression changes of DUXA and MLT1D family. To explore the potential roles of DUXA in bovine ZGA, we overlapped the genes impacted by siRNA knockdown with our newly defined ZGA gene list of IVT embryos. Furthermore, the expression levels of differentially expressed MLT1D loci were compared between DUXA knock down and control groups, and the significance of expression changes was calculated using Wilcoxon test.

## DATA AVAILABILITY

All datasets were collected from NCBI GEO database. RNA-seq of IVV embryo: GSE59186; RNA-seq of IVT embryo: GSE52415, GSE163620, GSE178438, GSE123705, GSE121227, GSE99294, GSE124100, GSE95311, GSE290265; RNA-seq of SCNT embryo: GSE178438, GSE123705, GSE121227, GSE125724, GSE99294, GSE124100, GSE114596, GSE99072, GSE95311. ATAC-seq of IVT embryo: GSE143658, GSE145040, GSE193640. CUT & Tag of IVT embryo: GSE193640. DUXA knockdown of IVT embryo: GSE264468. 5’end RNA-seq data of IVT embryo: GSE225056.

## ACKNOWLEDGEMENTS

The author thanks all members of the Duan lab and the IISAGE Consortium for the discussion of the TE analysis. This study was supported by grants from the NSF BII (#2213824) to J.E.D. and the IISAGE Consortium.

## AUTHOR CONTRIBUTIONS

G.L.: conceptualization, methodology, data curation, formal analysis, manuscript writing, and revision. C. F.: conceptualization, supervision, project administration, and manuscript revision. J. E. D.: conceptualization, supervision, project administration, manuscript revision, and funding acquisition. All co-authors participated in revising the manuscript.

## CODE AVAILABILITY

Scripts and analysis for genomic analysis are available at Github (https://github.com/Guangsheng357/Bovine-TE).

